# Similarities and distinctions in the activation of the *Candida glabrata* Pdr1 regulatory pathway by azole and non-azole drugs

**DOI:** 10.1101/2024.09.19.613905

**Authors:** Thomas P. Conway, Bao Gia Vu, Sarah R. Beattie, Damian J. Krysan, W. Scott Moye-Rowley

**Affiliations:** Departments of Molecular Physiology and Biophysics and Department of Pediatrics Carver, College of Medicine, University of Iowa, Iowa City, IA 52242 USA; Department of Microbiology and Immunology, University of Oklahoma Health Sciences Center, Oklahoma City, OK 73104

## Abstract

Incidences of fluconazole (FLC) resistance among *Candida glabrata* clinical isolates is a growing issue in clinics. The pleiotropic drug response (PDR) network in *C*. *glabrata* confers azole resistance and is defined primarily by the Zn^2^Cys_6_ zinc cluster-containing transcription factor Pdr1 and target genes such as *CDR1*, that encodes an ATP-binding cassette transporter protein thought to act as a FLC efflux pump. Mutations in the *PDR1* gene that render the transcription factor hyperactive are the most common cause of fluconazole resistance among clinical isolates. The phenothiazine class drug fluphenazine and a molecular derivative, CWHM-974, which both exhibit antifungal properties, have been shown to induce the expression of Cdr1 in *Candida* spp. We have used a firefly luciferase reporter gene driven by the *CDR1* promoter to demonstrate two distinct patterns of *CDR1* promoter activation kinetics: gradual promoter activation kinetics that occur in response to ergosterol limitations imposed by exposure to azole and polyene class antifungals and a robust and rapid *CDR1* induction occurring in response to the stress imposed by fluphenazines. We can attribute these different patterns of *CDR1* induction as proceeding through the promoter region of this gene since this is the only segment of the gene included in the luciferase reporter construct. Genetic analysis indicates that the signaling pathways responsible for phenothiazine and azole induction of *CDR1* overlap but are not identical. The short time course of phenothiazine induction suggests that these compounds may act more directly on the Pdr1 protein to stimulate its activity.

**Importance:** *Candida glabrata* has emerged as the second-leading cause of candidiasis due in part to its ability to acquire high level resistance to azole drugs, a major class of antifungal, that acts to block the biosynthesis of the fungal sterol ergosterol. The presence of azole drugs causes the induction of a variety of genes involved in controlling susceptibility to this drug class including drug transporters and ergosterol biosynthetic genes such as *ERG11*. We found that the presence of azole drugs leads to an induction of genes encoding drug transporters and *ERG11*, while exposure of *C. glabrata* cells to antifungals of the phenothiazine class of drugs caused a much faster and larger induction of drug transporters but not *ERG11*. Coupled with further genetic analyses of the effects of azole and phenothiazine drugs, our data indicate that these compounds are sensed and responded to differentially in the yeast cell.

## Introduction

Antifungal resistance among yeast pathogens continues to increase. The limited number of antifungal drugs primarily being used belong to three classes: azoles, polyenes, and echinocandins. Azole drugs are routinely used to treat fungal infections, with fluconazole (FLC) being among the most prescribed antifungal drug globally (1, 2). *Candida glabrata* is a human commensal and opportunistic fungal pathogen with a low intrinsic susceptibility to FLC and a high rate of developing increased resistance to azole drugs (Recently reviewed in (3)). Azoles, including FLC, inhibit function of the lanosterol α14-demethylase enzyme encoded by the *ERG11* gene, disrupting the ergosterol biosynthetic pathway and preventing fungal growth (4). The low FLC susceptibility of *C*. *glabrata* is attributed primarily to the functions of two Zn_2_Cys_6_ DNA-binding domain-containing transcription factors, Upc2A and Pdr1. Upc2A is a positive regulator of genes involved in ergosterol biosynthesis (*ERG* genes) and induces the expression of *ERG* genes in cells experiencing limited ergosterol availability, like that associated with azole stress (5). Pdr1 induces expression of the ATP-Binding Cassette (ABC) protein and putative drug efflux pump, Cdr1, as well as other genes in the pleiotropic drug response (PDR) pathway (6–8). Azole resistance among clinical isolates of *C. glabrata* is primarily due to nonsynonymous point mutations in the *PDR1* open reading frame which result in gain-of-function (GOF) Pdr1 isoforms. These GOF forms of Pdr1 cause constitutive high-level transcription of target genes with a corresponding decrease in FLC susceptibility (9). More recent data support a link between Upc2A and the Pdr system at the level of transcriptional control. Upc2A acts to coordinately induce expression of *ERG* genes with genes of the Pdr network, such as *PDR1* and *CDR1*, when ergosterol levels are reduced (10–12).

The limited number of antifungal agents has driven efforts to identify new therapeutic options for antifungal therapies. Among drugs that have been identified for their antifungal properties are those belonging to the phenothiazine molecular class (13). Fluphenazine (FPZ), a phenothiazine class antipsychotic medication, exhibits antifungal activity but its effective antifungal dosages exceed concentrations at which it can be safely used as a therapeutic agent (14). In *C*. *albicans*, FPZ induces the expression of ABC and Major Facilitator Superfamily (MFS) proteins associated with multi-drug resistance (15, 16). In 2018, Montoya et al tested FPZ derivatives and found that the analog CWHM-974 (called 974 here) has increased antifungal activity against *C*. *albicans* compared to FPZ (17). Miron-Ocampo et al (18) continued investigating the antifungal properties of FPZ and 974 in *Candida* species, demonstrating that both FPZ and 974 are potent inducers of *CDR1* in *C*. *albicans* and *C*. *glabrata*. It was also demonstrated that at subinhibitory concentrations the fluphenazine-derivatives antagonize FLC in *C*. *albicans*, but not in *C*. *glabrata* even though steady-state levels of Cdr1 protein were observed to increase by western blotting (18).

Here, we demonstrate that exposure of *C. glabrata* to either FPZ or 974 caused a strong and rapid transcriptional induction of *CDR1* mRNA. The fluphenazines induced higher levels of *CDR1* expression than FLC. In addition, *CDR1* was induced much more rapidly by the fluphenazines compared to FLC. Genetic analyses indicated that susceptibility of *C. glabrata* to these fluphenazine compounds responded to the level of Pdr1 activity. In contrast to the well-described effect of FLC on *ERG* gene expression, the phenothiazines did not significantly impact expression of genes in the ergosterol biosynthetic pathway. These data argue that activation of the Pdr1-*CDR1* pathway by azole and phenothiazine drugs occurs through both overlapping and distinct mechanisms.

## Materials/Methods

### Strains/media

*C. glabrata* strains were cultured at 30°C. Unless otherwise stated, cells were grown in YPD medium (1% yeast extract, 2% peptone, 2% glucose) for non-selective growth and drug treatment. For selective growth, cells were cultured in YPD supplemented with 50 µg/ml nourseothricin (NAT; Jena Bioscience, Jena, Germany) or complete synthetic medium (CSM) with appropriate amino acids omitted for heterotroph selection (Difco yeast nitrogen extract without amino acids, amino acid powder from Sunrise Science Products, 2% glucose). CSM media without methionine and supplemented with 1 mM estradiol was used to recycle the selection cassette associated with integration of different *PDR1* forms (19). All strains used in this study are listed in Table 1. CWHM-974 was synthesized as previously reported by the Meyers lab at St. Louis University (17).

**Table 1.** Strains used in this work.

| Name | Background | Genotype |
| --- | --- | --- |
| KKY2001 | CBS138 | *his3Δ::FRT leu2Δ::FRT trp1Δ::FRT* |
| SPG96 | KKY2001 | *his3Δ::FRT leu2Δ::FRT trp1Δ::FRT ura3Δ::FRT* |
| TCCG19 | KKY2001 | *HOΔ::CDR1-LUC::HIS3MX6* |
| TCCG92 | KKY2001 | *PDR1::loxP HOΔ::CDR1-LUC::HIS3MX6* |
| TCCG138 | KKY2001 | *pdr1Δ::loxP HOΔ::CDR1-LUC::HIS3MX6* |
| TCCG96 | KKY2001 | *R376W PDR1::loxP HOΔ::CDR1-LUC::HIS3MX6* |
| TCCG98 | KKY2001 | *D1082G PDR1::loxP HOΔ::CDR1-LUC::HIS3MX6* |
| TCCG203 | KKY2001 | *PDR1::loxP* |
| TCCG204 | KKY2001 | *R376W PDR1::loxP* |
| TCCG205 | KKY2001 | *D1082G PDR1::loxP* |
| TCCG16 | KKY2001 | *pdr1Δ::loxP* |
| TCCG110 | KKY2001 | *cdr1Δ::loxP* |
| TCCG52 | KKY2001 | *upc2AΔ::loxP* |
| TCCG123 | SPG96 | *med15AΔ::HIS3MX6* |
| TCCG51 | KKY2001 | *cna1Δ::loxP* |
| TCCG126 | KKY2001 | *crz1Δ::loxP* |
| TCCG125 | SPG96 | *bre5Δ::HIS3MX6* |
| TCCG206 | SPG96 | *ERG11::HIS3MX6* |
| TCCG207 | SPG96 | *Y141H,S410F ERG11::HIS3MX6* |
| TCCG51 | KKY2001 | *cna1Δ::loxP* |

### Luminescence Assay

For analysis of drug-induced *CDR1* promoter activation kinetics, a strategy for one-step measurement of firefly luciferase activity was employed (modified from (20). For this, a *CDR1* promoter-driven firefly luciferase reporter (*CDR1-LUC*) construct was integrated into the *HO* locus of *C*. *glabrata* strains analyzed (Figure 1A). The *CDR1-LUC* construct was flanked by regions of the HO gene (5’ and 3’) for integration. The *CDR1-LUC* fusion consisted of the entire intergenic region upstream of the *CDR1* start codon (−1 to −1734), placed upstream of the *Photinus pyralis* (Firefly) luciferase gene present in the plasmid pFA6-luc*(−SKL)-HIS3MX6 (Addgene #40233). (Figure 1A). For analyses of Pdr1-dependent *CDR1* promoter activation, isogenic Δ*pdr1*, wild-type *PDR1*, and two GOF forms of *PDR1* (R376W and D1082G) strains were derived from the *CDR1-LUC* parental strain using a *PDR1*-recyclable cassette (21) Strains were precultured overnight in YPD at 30°C and 200 rpm. The next morning, stationary phase cultures were diluted with fresh YPD to OD_600_ = 0.2. The diluted cultures were then grown at 30°C and 200 rpm until they reached mid-log phase growth (OD_600_ = 0.8).

**Figure 1.**
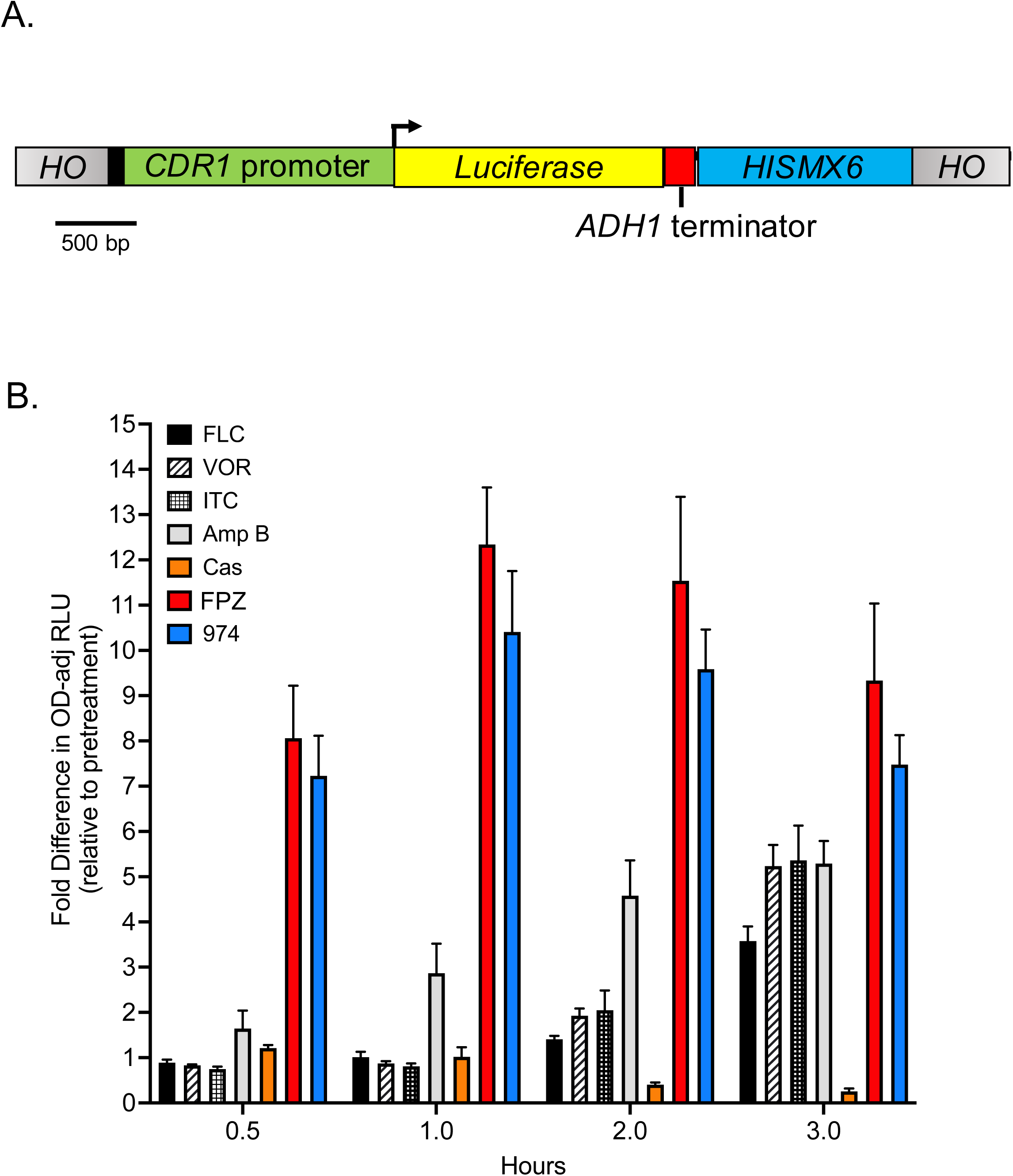
A comparison of drug-induced Cdr1 expression kinetics using a firefly luciferase reporter. (A) A luciferase reporter construct consisting of the firefly luciferase open reading frame immediately downstream of the full-length *CDR1* promoter was integrated into the *HO* locus of *Candida glabrata* strain KKY2001. (B) The reporter strain was grown to mid-log phase in rich liquid medium (YPD) and treated with drugs known to have antifungal properties. These experiments were performed in a 96-well format. Each drug was administered at its minimum inhibitor concentration (MIC). Drugs tested included: fluconazole (FLC; 16 μg/ml), voriconazole (VOR; 0.25 μg/ml), itraconazole (ITR; 0.5 μg/ml), amphotericin B (AMB; 0.25 μg/ml), caspofungin (CAS; 1 μg/ml), fluphenazine (FPZ; 32 μg/ml), and the FPZ analog CWHM-974 (974; 4 μg/ml). At 0.5, 1, 2, and 3 hours post treatment, optical density of each culture was measured at 600nm (OD_600_) and luciferase activity was measured in relative light units (RLU) after the addition of D-luciferin. Here, we report the OD-adjusted RLU of treated cells relative to their pretreatment state, which serves as a surrogate for fold induction of Cdr1 expression. Bars indicate mean of 4 replicates and error bars indicate SD.

Then, 50 µl of cell culture was pipetted into wells of a 96-well plate that contained either 50 µl of untreated YPD or 50 µl of YPD containing a drug at double its minimum inhibitory concentration (MIC). Thus, at the onset of the experiment each well contained 100 µl of culture at an OD_600_ = 0.4. For wells containing drug, the final concentration was equal to the MIC of the corresponding drug. During the experiment, plates were incubated at 30°C without shaking.

At each time point analyzed for luciferase expression, the OD_600_ and luminescence measurements were acquired in duplicate from two independent wells for each condition. Prior to the addition of D-luciferin substrate to wells, OD_600_ was measured using a SpectraMax iD3 plate reader (Molecular Devices, San Jose, CA) set to measure absorbance (ABS) at wavelength 600. Subsequently, 100 µl of 1 mM D-luciferin potassium salt (Perkin Elmer, Waltham, MA) in 0.1M sodium citrate buffer (pH 5) was pipetted into the appropriate culture wells. The plate was then immediately measured for luminescence at all wavelengths using a SpectraMax iD3 plate reader with integration time set to one second for each well. The luminescence measurements are expressed as relative light units (RLU) and were normalized by dividing the luminescence value given by the luminometer by the well-specific OD_600_, which yielded the OD-adjusted RLU measurements used for calculating the fold induction. Data represented in the associated graphs are the average of two independent biological replicates each with a minimum of two technical replicates.

### Spot Dilution Assay

*C*. *glabrata* strains were grown to mid-log phase and spotted in ten-fold serial dilutions on YPD agar plates containing the indicated concentrations of fluconazole (LKT Laboratories), fluphenazine (Sigma-Aldrich), or CMHW-974 (17, (18). Plates were incubated at 30°C for 24 to 48 hours prior to imaging.

### RT-qPCR assay

To analyze transcriptional activation of azole-induced genes in wild-type and deletion mutants, samples containing six OD_600_ units of mid-log phase cells were acquired prior to drug exposure and the specified time points after addition of drugs to the cultures. Total RNA was extracted from cell samples using Trizole (Invitrogen #15596026) and chloroform. For RNA purification, a RNeasy Mini Kit (Qiagen #74104) was used, and cDNA was generated from 0.5 µg of purified RNA using an iScript cDNA Synthesis Kit (Bio-Rad #1708890). iTaq Universal SYBR Green Supermix (Bio-Rad #1725151) was used for qPCR, and transcript levels of target genes were normalized to 18s rRNA transcript levels. The ΔΔCT method was used to calculate fold change in transcript levels between pretreatment and treatment samples. All data presented are the averaged result of two biological replicates each with two technical replicates, for a total of four replicates. Statistical analysis was performed using a one-way ANOVA with Tukey’s or Dunnett’s multiple comparison test.

## RESULTS

### The kinetics of *CDR1* induction by fluphenazines are distinct from other antifungal drugs

Earlier work demonstrated that exposure of *C. glabrata* cells to either FPZ or 974 led to the rapid increase in levels of the Cdr1 ABC transporter protein as measured by western blotting with anti-Cdr1 antiserum (18, 22). Analyses of FLC induction of *CDR1*, both at the transcription and protein level (22), indicated that the response to this drug required a longer period of drug exposure to see full induction. To facilitate comparison of FLC, FPZ, and 974-mediated induction of *CDR1*, we prepared a translational fusion between a firefly luciferase (*LUC*) gene and the *CDR1* promoter. This *CDR1-LUC* fusion gene was then integrated into the *C. glabrata* genome at the *HO* locus (Figure 1A). This reporter gene allowed use of a 96-well format for rapid assay of *CDR1* promoter activation over time as well as with a variety of different antifungal drugs. We used this strain containing the *CDR1-LUC* fusion gene to compare the induction time courses for three different azole drugs (FLC; voriconazole, VOR; itraconazole, ITC), amphotericin B (AmB), caspofungin (CAS) as well as the two phenothiazine derivatives. The levels of *CDR1*-driven luciferase activities were measured for all these different conditions and are shown in Figure 1B.

Both fluphenazines triggered a very rapid and large (∼8-fold) induction of *CDR1-LUC* expression after only 30 minutes of exposure. AmB induced 3-fold *CDR1-LUC* activity after 1 hour with this level of induction plateauing at 5-fold after 3 hours of AmB treatment. The azole drugs required almost 3 hours of exposure before reaching a similar induction level to that seen for AmB. CAS exposure did not lead to any significant changes in *CDR1* expression in this assay.

While these effects on the *CDR1-LUC* fusion gene were provocative, we wanted to ensure that the native *CDR1* gene also exhibited the rapid induction kinetics seen for the reporter gene. Additionally, we tested expression of the *PDR1* gene that is known to be autoregulated (23) and *CDR2* as an additional Pdr1 target gene (8). Expression of these three genes was assessed using RT-qPCR to measure steady-state mRNA after exposure to FLC or the two phenothiazine drugs (Figure 2).

**Figure 2.**
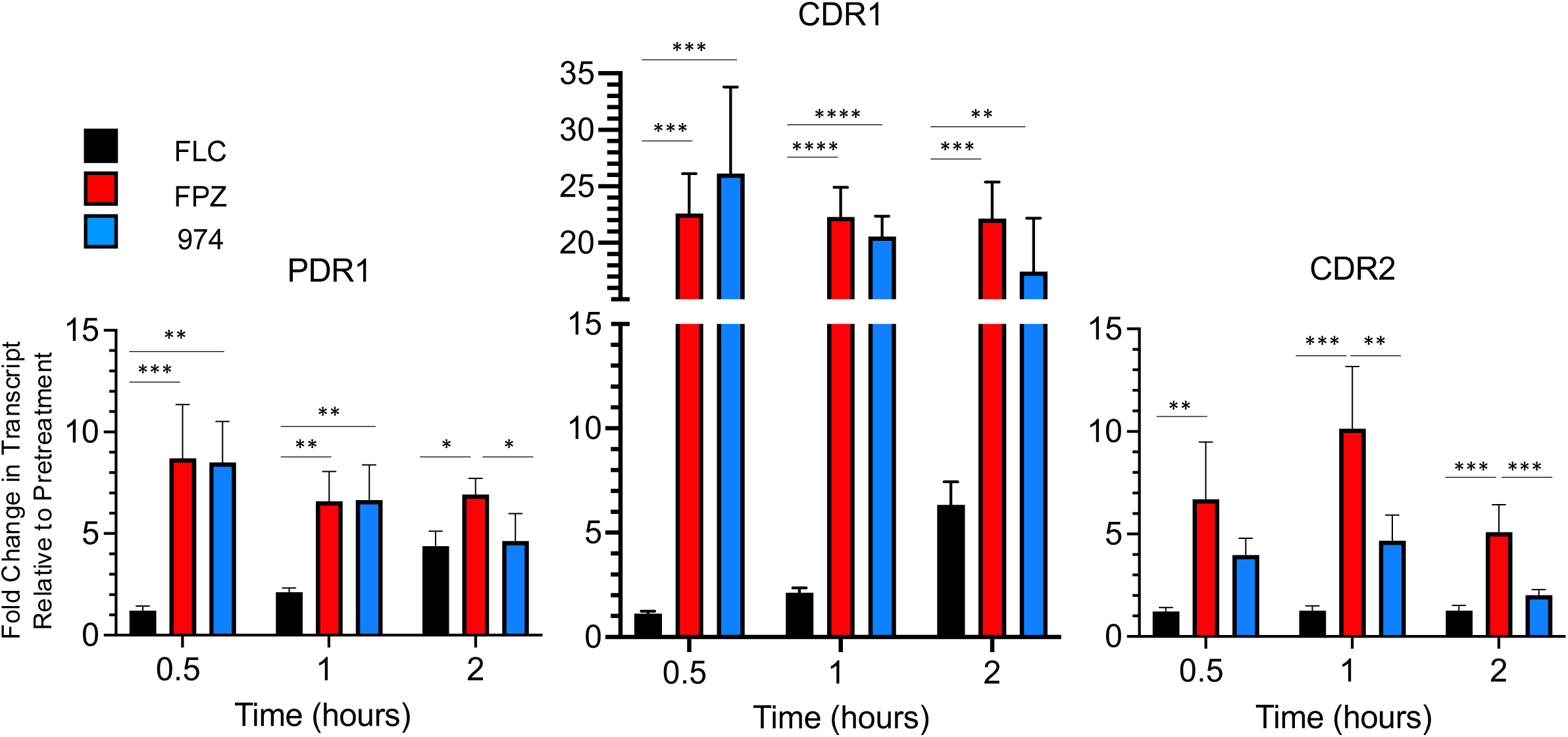
Phenothiazines show differential activation of *PDR1*, *CDR1*, and *CDR2* transcription compared to fluconazole. Mid-log phase *C. glabrata* cells were treated with fluconazole (FLC), fluphenazine (FPZ), or CWHM-974 (974) at their minimum inhibitory concentrations (16 μg/ml, 32 μg/ml, and 4 μg/ml, respectively). Samples were acquired pretreatment and 0.5, 1, and 2 hours post treatment for analysis of transcriptional changes that occurred for *PDR1* (A), *CDR1* (B), and *CDR2* (C) in response to each drug. Data is represented as fold change in transcript levels compared to pretreatment. Each data point is the average of two biological replicates each with two technical replicates, and a one-way ANOVA with Tukey’s multiple comparison test was used for statistical analyses. Significance is displayed as: *P<0.05, **P<0.01, ***P<0.001, ****P<0.0001.

The wild-type *CDR1* gene was rapidly induced upon exposure to either phenothiazine, rising to more than 20-fold compared to pre-treatment transcript levels after only 30 minutes of drug challenge. Both the rate and magnitude of induction after phenothiazine treatment exceeded that seen for FLC exposure which required 2 hours to cause a 6-fold increase in *CDR1* transcript levels. Similar rapid induction kinetics have been reported for *CDR1* mRNA recently (24). Similarly, both *PDR1* and *CDR2* were rapidly induced by phenothiazine treatment. *PDR1* mRNA was increased by 5-fold during FLC treatment but, as we have seen before (25), *CDR2* transcript levels were not altered by the presence of FLC.

### Resistance to phenothiazines is Pdr1-dependent

The strikingly different *PDR1*/*CDR1* induction kinetics of the phenothiazines compared to FLC prompted us to test the effect of *PDR1* mutants on phenothiazine susceptibility. As mentioned above, Pdr1 is required for FLC induction of *CDR1* and the most common causes of FLC resistance in *C. glabrata* clinical isolates are gain-of-function (GOF) mutants of Pdr1. As previously described, we prepared isogenic *pdr1Δ* strains along with two different GOF forms of *PDR1*: R367W and D1082G (9, 26). These strains were then tested for their ability to grow on rich media containing various concentrations of FLC or the two phenothiazine drugs (Figure 3A).

**Figure 3.**
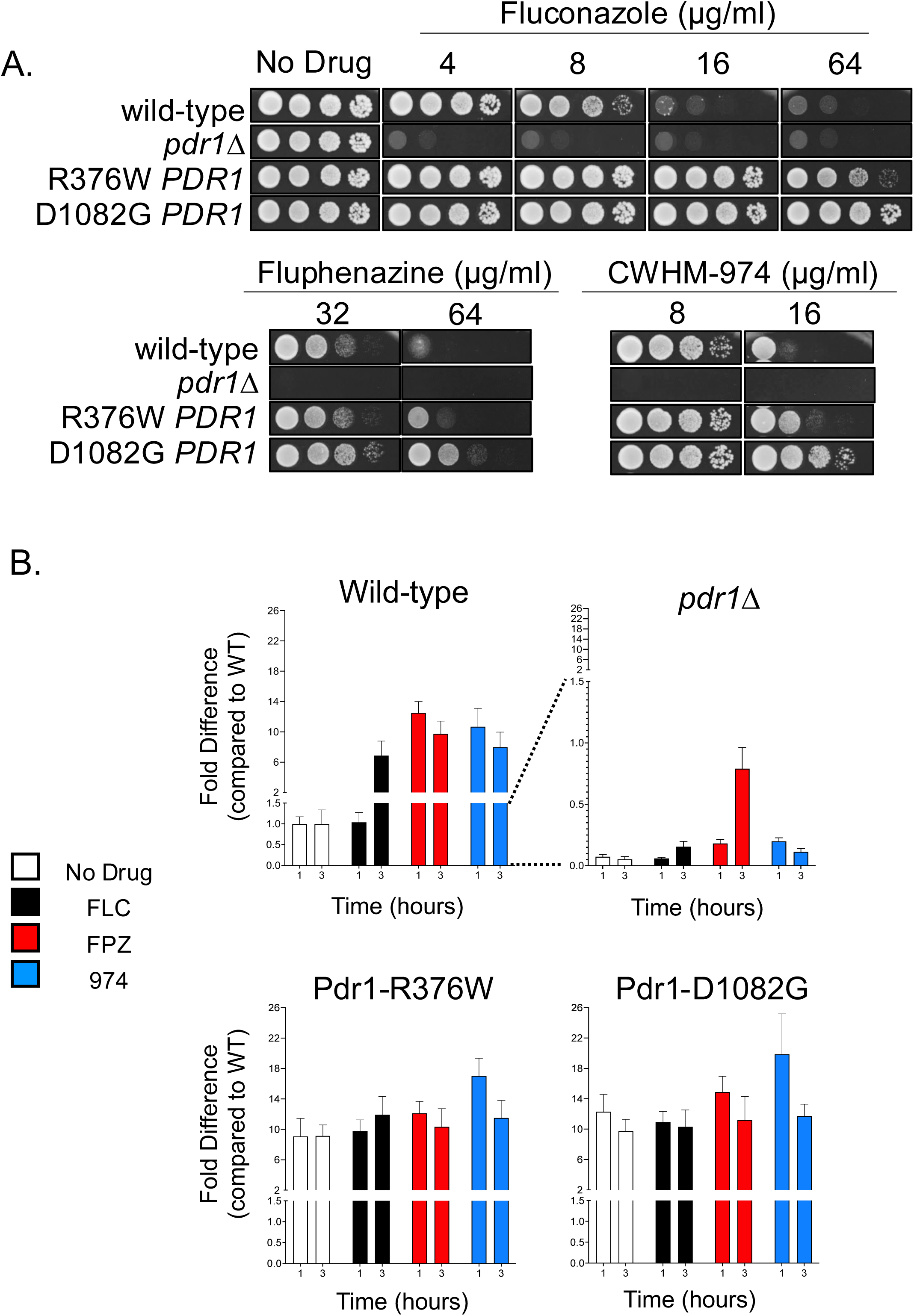
Phenothiazine resistance correlates with Pdr1 activity and the expression of Cdr1. (A) A spot test assay was used for analysis of phenothiazine resistance of *Candida glabrata* strains varying only at their *PDR1* locus. Strains expressing wild-type Pdr1, gain-of-function forms of Pdr1 (Pdr1-R376W or Pdr1-D1082G), or in which the *PDR1* open reading frame was deleted (*pdr1Δ*) were grown to mid-log phase and serial dilutions were spotted on rich agar medium (YPD) contain varying concentrations of fluconazole (FLC), fluphenazine (FPZ), or CWHM-974 (974). (B) Strains expressing wild-type Pdr1, R376W-Pdr1, D1082G-Pdr1, or lacking a *PDR1* open reading frame (*pdr1Δ*) were modified to express firefly luciferase under the *CDR1* promoter. Each strain was grown in the absence of drug (No Drug) or in the presence of minimum inhibitory concentrations of FLC (16 μg/ml), FPZ (32 μg/ml), or 974 (4 μg/ml). Luciferase expression was measured as described in Materials and Methods at one and three hours post treatment and compared to pretreatment levels of the strain expressing wild-type Pdr1.

The susceptibility of the *PDR1* mutants to phenothiazines was very similar to the patterns previously observed for fluconazole. Specifically, loss of *PDR1* increased susceptibility to both classes of drugs while the GOF Pdr1 mutants caused a significant decrease in drug susceptibility. Interestingly, the D1082G form of Pdr1 exhibited a greater decrease in phenothiazine susceptibility compared to the R376W Pdr1 protein as we previously reported for FLC susceptibility (26).

We also integrated the *CDR1-LUC* reporter gene into these strains and compared azole- and phenothiazine-induced *CDR1* activation (Figure 3B). The presence of *PDR1* was required for wild-type induction by all drugs tested. A small degree of FPZ induction was observed after 3 hours in the *pdr1Δ* strain but this remained at only 10% of that seen in the isogenic wild-type background. Both GOF forms of Pdr1 drove high, constitutive levels of *CDR1-LUC* that was not further increased by drug exposure. As previously seen for FLC, the *PDR1* gene is a key determinant of both phenothiazine-induced *CDR1* expression and susceptibility

### Genes impacting Pdr1-mediated FLC susceptibility have similar but not identical effects on phenothiazine susceptibility

Having confirmed a Pdr1-dependent mechanism for phenothiazine-induced *CDR1* expression and resistance, we examined the contribution of genes previously identified as being involved in the *PDR1*-mediated FLC resistance pathway. We used strains that lacked *CDR1*, *PDR1*, or a number of different proteins that have been implicated in Pdr1-mediated FLC susceptibility.

*UPC2A* is a transcription factor required for upregulation of genes involved in ergosterol biosynthesis and functions together with Pdr1 in azole-induced activation of Cdr1 expression (11, 27). *MED15A*, a nonessential subunit in the tail of the Mediator complex, has been demonstrated to interact directly with Pdr1 and is required for Pdr1-directed gene activation and azole resistance (28). Med15 was also found to be required for Tac1-dependent FPZ induction of *CDR1* in *C. albicans* (29). *BRE5* encodes a protein subunit of a deubiquitinase complex and interacts with Pdr1 as a negative regulator (30). *CNA1* encodes the catalytic subunit of the protein phosphatase calcineurin that we showed is a positive regulator of Pdr1 (21). In addition to the role of Cna1 in fluconazole-induced gene expression, it is also notable that a deletion mutant of calcineurin is hypersusceptible to FPZ in *Candida* species (31). Accordingly, *CRZ1* was included in this screen as it encodes the stress-responsive transcription factor that is an important protein target of calcineurin (reviewed in (32)). To compare the role of the above set of genes in azole and phenothiazine susceptibility, we used a spot test assay to analyze the phenotype of single gene deletion mutants grown on a rich medium with varying concentrations of FLC, FPZ, and 974.

Consistent with previous observations, the individual deletion mutants of *CDR1*, *PDR1*, *UPC2A*, *MED15A*, or *CNA1* increased susceptibility to FLC, and the deletion of *BRE5* decreased FLC susceptibility (Figure 4A). Surprisingly, the deletion of *CRZ1* also resulted in decreased fluconazole susceptibility comparable to that observed for the *bre5Δ* mutant (Figure 4A). On plates containing FPZ or 974, no deletion mutant analyzed exhibited decreased susceptibility compared to the wild-type control. The *pdr1Δ* and *cna1Δ* strains exhibited the highest level of susceptibility to the phenothiazines (Figure 4B,4C). Deletion mutants of *cdr1Δ*, *pdr1Δ*, *med15AΔ*, and *cna1Δ* were unable to grow on media containing 48 µg/ml FPZ while deletion of *UPC2A*, *BRE5*, or *CRZ1* had relatively minor effects (Figure 4B). In the presence of 8 μg/ml 974, loss of *CDR1*, *BRE5*, and *CRZ1* had minor effects while loss of *UPC2A* or *MED15A* produced a strain nearly as susceptible as the *pdr1Δ* or *cna1Δ* strains (Figure 4C). While there was overlap in the genes involved in FLC and phenothiazine susceptibility, significant differences in the response to loss of particular regulators emerged. Loss of *CDR1*, *PDR1*, *UPC2A* and *MED15A* caused profound FLC sensitivity while a *cna1Δ* strain was more susceptible to phenothiazines than these mutants.

**Figure 4.**
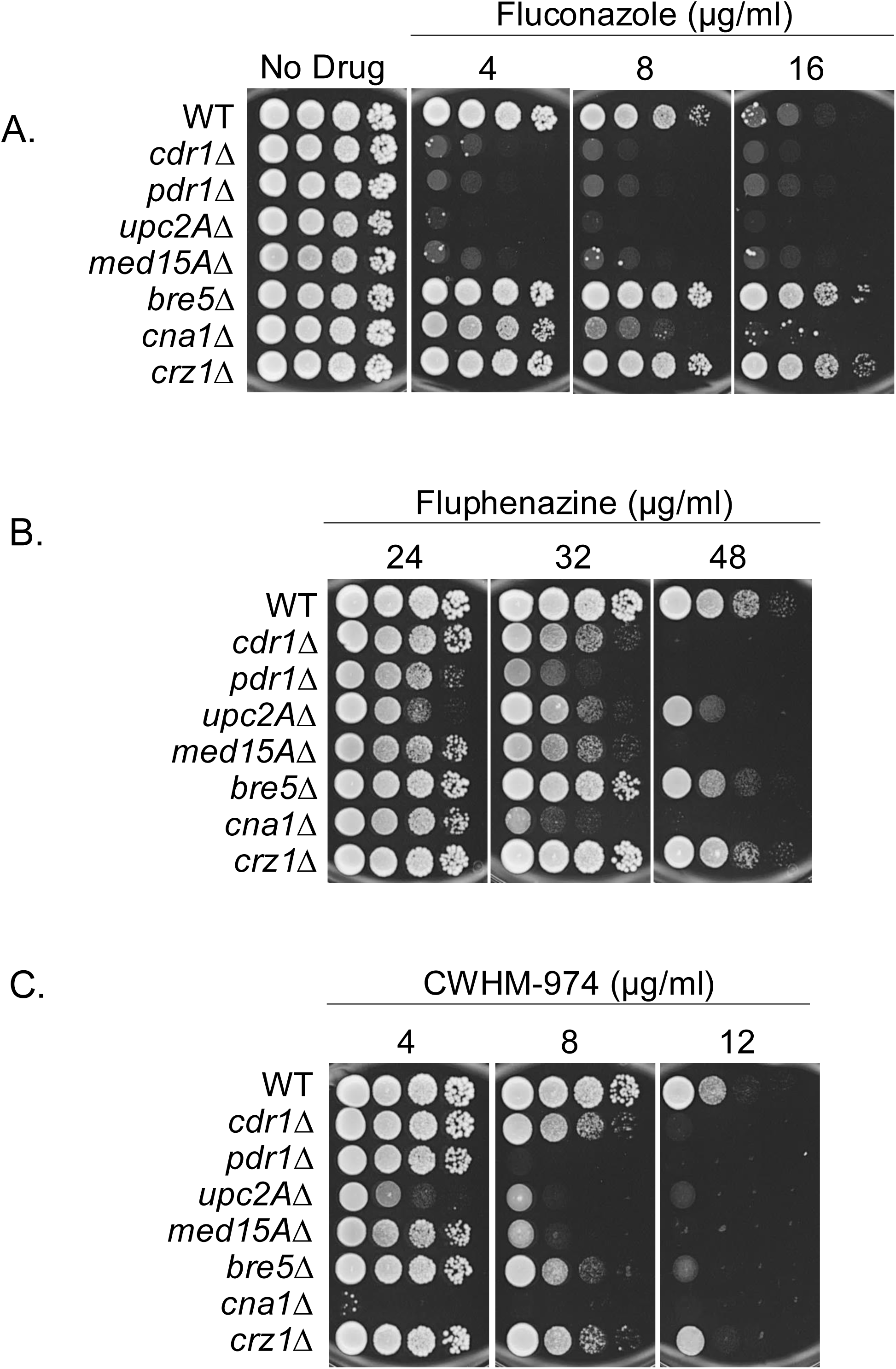
Single gene deletions affecting azole resistance differentially affect resistance to fluphenazine and CWHM-974. Single gene deletions that affect fluconazole resistance in *Candida glabrata* (i.e. *cdr1Δ, pdr1Δ, upc2AΔ, med15AΔ, bre5Δ, cna1Δ, and crz1Δ*) were tested for altered resistance to fluphenazine and CWHM-974. Strains were grown to mid-log phase and serial dilutions were spotted on YPD containing fluconazole (A), fluphenazine (B), or CWHM-974 (C) at varying concentrations for comparison of susceptibility to each antifungal compound. Plates were imaged and susceptibility phenotypes assessed after 48 hours incubation at 30°C.

To correlate these genetic differences in drug susceptibility with effects on gene expression, we analyzed the transcription of *PDR1, CDR1, and ERG11,* three genes important in FLC susceptibility, using RT-qPCR The strains described above were grown to mid-log phase, treated with FLC, FPZ or 974 for 2 hours and total RNA prepared.

Drug-induced expression of *PDR1* was reduced in the absence of *MED15A*, *CNA1*, or *CRZ1*, reaching significance in 7 out of 9 conditions (Figure 5A). The magnitude of *PDR1* expression induced by the phenothiazines was the same as that induced by FLC (A maximum of approximately 4-fold in response to all treatments). The relative effects of FLC and phenothiazines on *CDR1* were quite different by comparison with *PDR1* transcription (Figure 5B). *CDR1* induction with FLC did not exceed 8-fold while induction with the phenothiazines ranged from 15- to 37-fold. For both FPZ and 974, loss of either *PDR1* or *MED15A* blocked induction while induction by 974 was reduced in the absence of *UPC2A, ERG11* expression was essentially unaffected by the phenothiazine drugs (Figure 5C). *ERG11* mRNA was induced by FLC by at least 2-fold in all mutants tested with the exception of the *upc2AΔ* strain that exhibited the expected reduction in *ERG11* expression.

**Figure 5.**
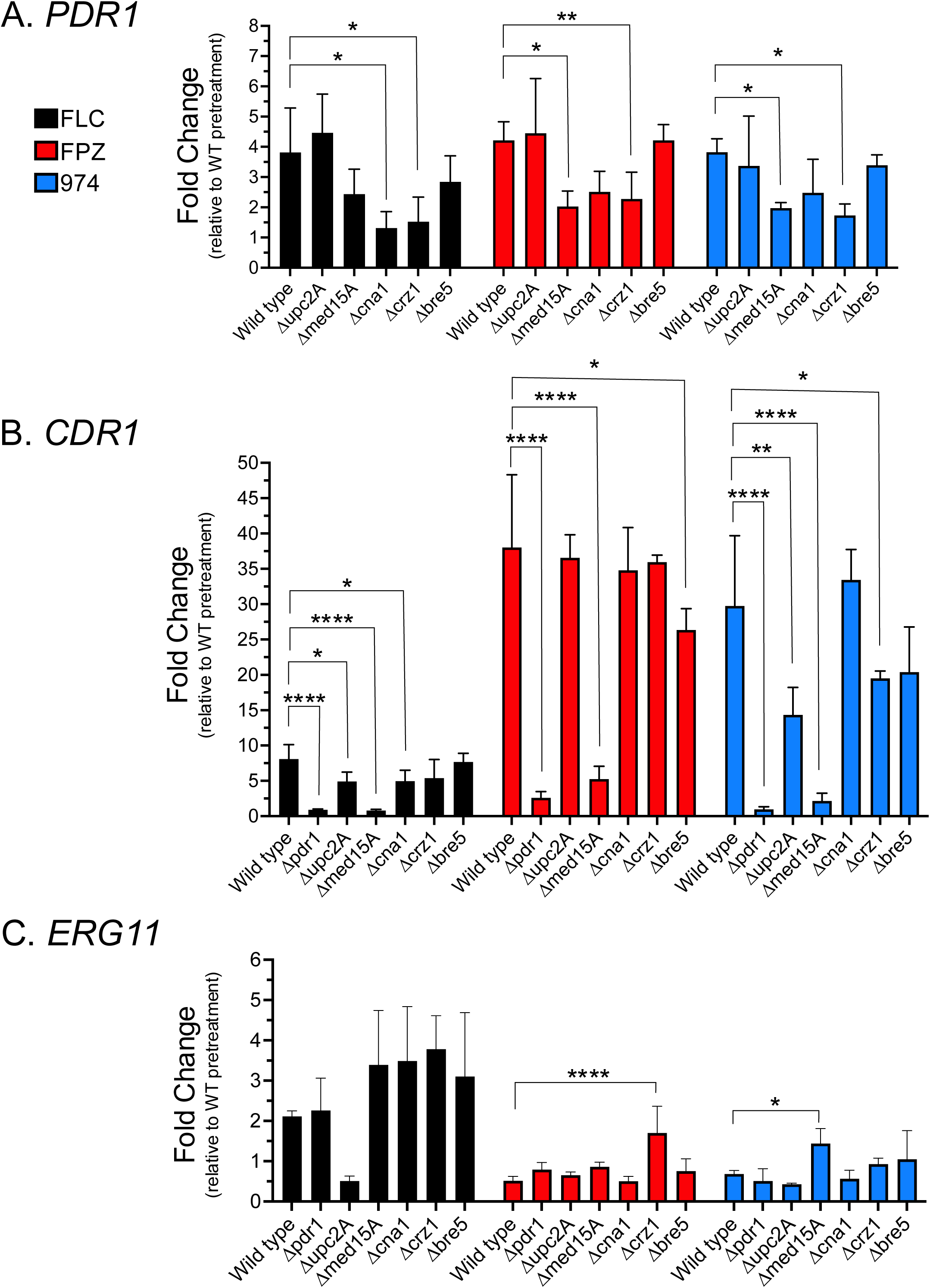
Analysis of the effect of single gene deletions on activated transcription of *PDR1*, *CDR1* and *ERG11* in response fluconazole, fluphenazine, and CWHM-974. Single gene deletions that affect fluconazole resistance in *Candida glabrata* (i.e. *cdr1Δ*, *pdr1Δ*, *upc2AΔ*, *med15AΔ*, *bre5Δ, cna1Δ*, and *crz1Δ*) were analyzed for effects on induced transcription of *PDR1* (A), *CDR1* (B) and *ERG11* (C). *C. glabrata* strains were cultured to mid-log phase in liquid YPD and split between three conditions: (1) fluconazole (FLC; 16 μg/ml), (2) fluphenazine (FPZ; 32 μg/ml), and CWHM-974 (974; 4 μg/ml). Samples were acquired pretreatment and two hours post treatment for each condition tested. RT-qPCR was used for analysis of changes in transcript levels. Data is displayed as fold change in transcript levels for each gene analyzed relative to pretreatment levels of the wild-type control strain. Each data point is the average of two biological replicates each with two technical replicates, and a one-way ANOVA with Dunnett’s multiple comparison test was used for statistical analyses. Significance is displayed as: *P<0.05, **P<0.01, ***P<0.001, ****P<0.0001.

### The phenothiazines do not affect the *ERG* pathway

Previous work indicated that the presence of FLC or mutant forms of Erg11 activates both the *PDR* and *ERG* pathways (11, 12, 22). Accordingly, one potential mechanism by which the phenothiazines could contribute to *CDR1* expression is by interference with ergosterol biosynthesis. If that were operative, then we would expect that *ERG* gene expression would be increased in FPZ/974-treated cells as is the case for FLC. Therefore, we more closely examined the effects of the phenothiazines on a second *ERG* pathway gene.

First, we used RT-qPCR to examine the effect of phenothiazines on the transcription of both *ERG11* and a gene acting later in the ERG pathway (*ERG4*) (Figure 6A). FLC induced both *ERG11* and *ERG4* transcription at both 1& 2 hour after treatment while neither phenothiazine affected *ERG* gene expression at those time points.

**Figure 6.**
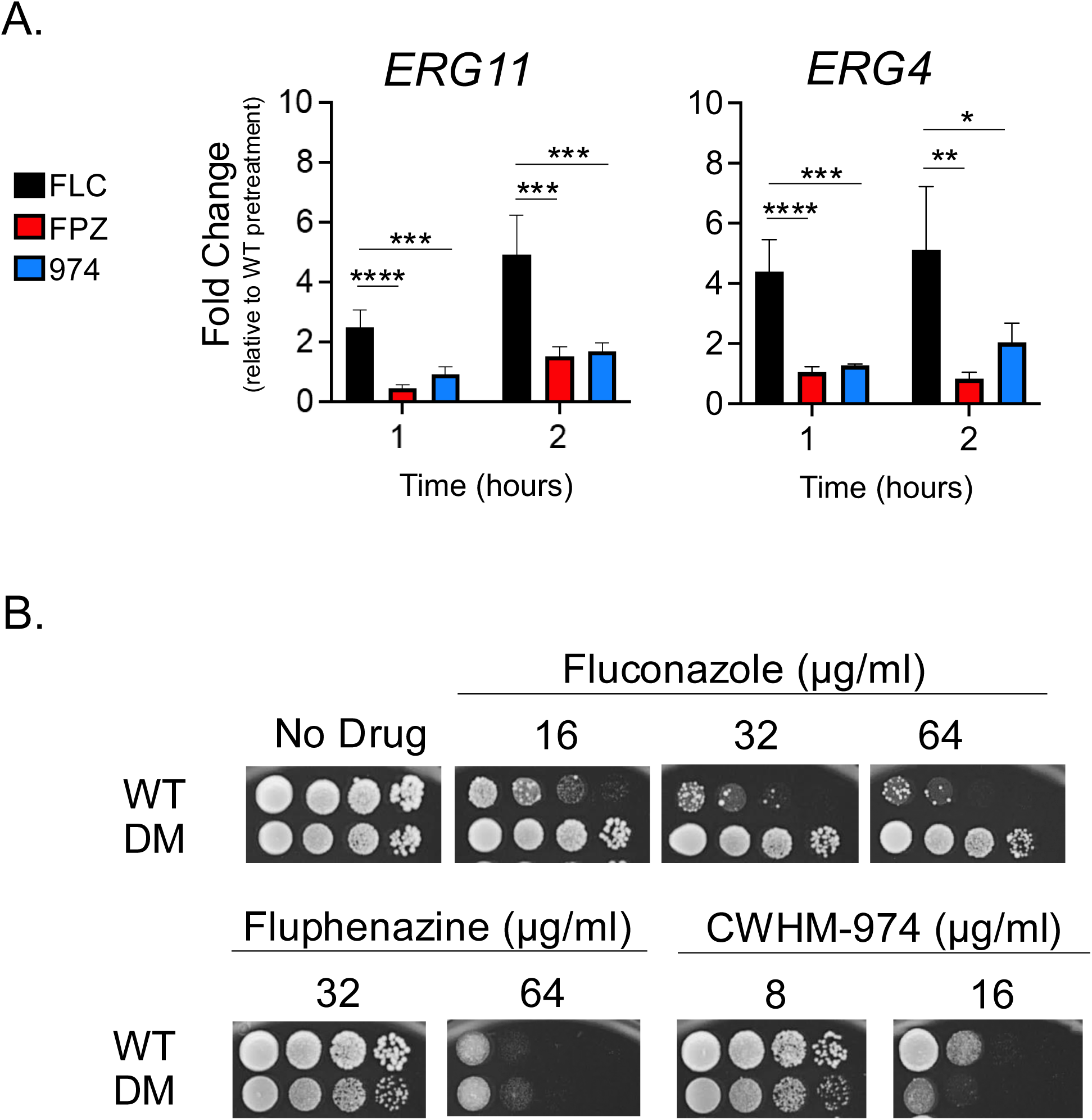
Acute effects on the ergosterol biosynthetic pathway imposed by exposure to azoles and phenothiazines are nonidentical. (A-B) Transcript levels of *ERG11* (A) and *ERG4* (B), each a gene encoding an enzyme with a different role in the ergosterol biosynthesis pathway, were examined one and two hours after exposure to the minimum inhibitory concentrations (MICs) of fluconazole (FLC; 16 μg/ml), fluphenazine (FPZ; 32 μg/ml), or CWHM-974 (974; 4 μg/ml). Data is represented as fold change in transcript levels relative to pretreatment levels. (D) To further investigate the differential impact of FLC, FPZ, and 974 on the ergosterol biosynthesis pathway we examined the susceptibility of an *ERG11* double mutant (DM; Erg11-Y141H,S410F) previously documented as hyper-resistant to fluconazole (12). The wild-type (WT) and DM strain were grown to mid-log phase and serial dilutions were spotted on plates containing varying concentrations of FLC, FPZ, or CWHM-974. Plates were imaged and susceptibility phenotypes assessed after 48 hours incubation at 30°C. Each data point is the average of two biological replicates each with two technical replicates, and a one-way ANOVA with Tukey’s multiple comparison test was used for statistical analyses. Significance is displayed as: *P<0.05, **P<0.01, ***P<0.001, ****P<0.0001.

Second, we determined the phenothiazine susceptibility of a strain containing a double mutant form of *ERG11* (Y141H and S410F) that we have previously shown to cause a strong decrease in FLC susceptibility. Isogenic wild-type and Y141H S410F *ERG11* strains were grown to mid-log and then tested by spotting dilutions on varying concentrations of FLC, FPZ, and 974 (Figure 6B). Consistent with the lack of a phenothiazine effect on *ERG* gene expression, the Y141H S410F *ERG11* strain exhibited negligible effects on susceptibility of these drugs while causing the previously reported strong decrease in FLC susceptibility. These data indicate that the phenothiazines do not affect *ERG* gene expression and are unlikely to interfere with ergosterol biosynthesis through other mechanisms. These observations further support the distinct nature of phenothiazine action versus that of FLC even though both drug classes trigger activation of *CDR1* expression.

## Discussion

Our finding of the differential kinetics and magnitude of *C. glabrata CDR1* induction by phenothiazine drugs compared to FLC prompted our investigation of the molecular basis of this difference. FLC (and other azole drugs) led to a relatively slow but steady increase in *CDR1* expression over the same time course during which both phenothiazines triggered a rapid and much larger induction (Figure 1). Here we demonstrate that the effects of the phenothiazines occurs at the level of *CDR1* transcription and involves many of the same regulatory factors as previously required for FLC induction. However, the fluphenazines trigger both a faster and larger induction of some Pdr1-regulated genes than does FLC. There are multiple additional distinctions between FLC and fluphenazine-induction of Pdr1 regulated genes. First, our previous experiments demonstrated that *CDR2* is not responsive to FLC challenge (25) but here is induced by 5-fold or more by phenothiazine treatment. Second, the autoregulation of *PDR1* (as measured by induction of *PDR1* mRNA levels) was more rapid with the phenothiazines but FLC-induced autoregulation eventually reached very similar levels, although with a slower time course.

There are at least two different explanations for the rapid induction of the three different Pdr1 target genes especially for *PDR1* itself. Treatment of cells with the phenothiazine drugs led to a large increase in *PDR1* mRNA within 30 minutes of exposure (Figure 2). This is an extremely rapid time course and much faster than the induction seen in the presence of FLC. We believe the most likely explanation for this difference is a more direct action of the phenothiazines on Pdr1 itself. Since the *PDR1* gene is autoregulated (23), direct stimulation of the function of Pdr1 will trigger both the increase of Pdr1 protein levels as well as *PDR1* mRNA. Increased Pdr1 transcription factor activity would be sufficient to explain the observed increase in *CDR1* and *CDR2* expression since both of these genes respond to changes in Pdr1 activity (8, 33). A second possibility is the presence of some other factor that can both be activated by phenothiazine exposure and also regulates *PDR1*, *CDR1*, and *CDR2* transcription. Further experiments are required to discriminate between these different modes of gene activation for these Pdr1 target genes.

AmB treatment of cells led to a faster induction of *CDR1-LUC* compared to the azole drugs but after 3 hours, levels of *CDR1* expression were equivalent across these different conditions. AmB directly binds to ergosterol in the plasma membrane while the azole drugs cause ergosterol depletion by inhibiting biosynthesis of this essential membrane lipid (recently reviewed in (34)). The faster induction caused by AmB exposure suggests that direct alteration of membrane ergosterol levels may lead the generation of a more proximal signal causing *CDR1* activation. The AmB induction of *CDR1* expression could explain the observed antagonism between this polyene drug and azole antifungals (35) in certain situations.

A common theme of all these signals that induce *CDR1* promoter is the sufficiency of the promoter region to explain the observed effect. This is one of the advantages of using the *CDR1-LUC* reporter system as the only segment of the *CDR1* that is present is the 1.7 kb 5’ noncoding sequence of the *CDR1* locus. Our previous demonstration of phenothiazine induction of *CDR1* relied on the use of anti-Cdr1 antiserum to determine that steady-state levels of Cdr1 were rapidly induced upon treatment with these compounds (18). This increase in Cdr1 could occur at multiple different levels in the context of the native *CDR1* gene but our current assays using the *CDR1-LUC* reporter system indicate an approximately 10-fold induction in luciferase levels upon phenothiazine treatment that agree well with the previous western blot data (18).

Phenothiazines have been used to induce and study *CDR1* transcription in *C. albicans* for some time as FLC was not thought to significantly activate *CDR1* transcription (36). FLC is well-known to strongly induce *CDR1* transcription in *C. glabrata* and the work shown here establishes that the phenothiazines also activate *CDR1* gene expression. Unlike *C. albicans*, neither phenothiazine was antagonistic to FLC in *C. glabrata* cells. FPZ increased the FLC MIC while 974 did not in assays of *C. albicans* drug susceptibility (18). This differential behavior of FPZ and 974 in *C. albicans* compared to *C. glabrata*, even though both phenothiazines induce *CDR1* transcription in these yeasts, illustrates the different downstream impacts of these drugs. We tested a collection of mutant strains that have known impacts on FLC regulation of *CDR1* and found that there was relatively poor correlation between FLC and phenothiazine susceptibilities for these strains. The largest effects on phenothiazine susceptibilities were seen for strains lacking *PDR1*, *CNA1* and *MED15A*. Loss of *CDR1* increased phenothiazine susceptibility but this was best seen at higher drug dosages.

These data argue that phenothiazine induction of *CDR1* expression occurs in a very different manner than FLC induction. FLC induction of *CDR1* proceeds in a Cna1-dependent manner as we have shown before (21). In contrast, loss of *CNA1* had a negligible effect on *CDR1* induction by phenothiazines (Figure 5B). The action of phenothiazines to inhibit calmodulin function (and subsequently calcineurin) would be expected to block Pdr1 activation and *CDR1* induction. The profound induction of Pdr1 by phenothiazine argues that the effect of these compounds on activity of this transcription factor cannot be explained by their calcineurin inhibition. We believe that FLC activation of Pdr1 proceeds in a calcineurin-dependent manner while phenothiazine stimulation of Pdr1 is calcineurin-independent and may be due to more direct interaction between these drugs and Pdr1. This would also explain the rapid time course seen for *CDR1* induction following phenothiazine treatment of *C. glabrata* cells.

The level of Pdr1 activity is an important determinant of phenothiazine susceptibility as can be illustrated by the analysis of GOF *PDR1* alleles (Figure 3). Both GOF forms of Pdr1 reduce phenothiazine susceptibility while loss of *PDR1* increases susceptibility to phenothiazines. We have previously demonstrated that the D1082G *PDR1* allele has a more prominent effect on FLC expression and susceptibility than the R376W Pdr1 allele (26) and this same behavior is also observed with respect to phenothiazine susceptibility.

We interpret these data to indicate that Pdr1-dependent transcriptional activation is an important component of the response to phenothiazine exposure. The *CDR1* gene does contribute to phenothiazine susceptibility but, unlike its central role in FLC resistance, this contribution is reduced. This can be appreciated by comparing the lack of growth of *cdr1Δ* cells on FLC at the lowest concentration tested (Figure 4A) while this same strain is only slightly reduced in growth on the phenothiazines (Figure 4B and 4C). We suggest that Pdr1 has target genes in addition to *CDR1* that are required for wild-type susceptibility to phenothiazines. *C. albicans* strains lacking *CDR1* also showed no increased susceptibility to phenothiazines (18).

Although phenothiazines are likely to have multiple targets, their ability to inhibit calmodulin in eukaryotic cells is well established and likely contributes to their antifungal activity. Phenothiazines are thought to cause toxicity primarily by inhibiting the calcium-binding regulatory protein calmodulin (17). Loss of calcineurin (*CNA1*) dramatically sensitizes cells to the phenothiazines yet deletion of the Cna1 target transcription factor Crz1 had no effect on phenothiazine susceptibility. While calmodulin is a well-established activator of calcineurin activity, these data establish that phenothiazine toxicity is not caused by inhibition of calcineurin activation of Crz1 in *C. glabrata*. As calmodulin has many targets in cells beyond calcineurin, loss of *CNA1* may block a compensatory response triggered by phenothiazines inhibition of calmodulin. Loss of Pdr1 could lead to increased phenothiazine levels in cells or cause some other defective response that prevents normal phenothiazine susceptibility.

## Acknowledgements

This work was supported by NIH grants AI152494 (WSM-R), AI168509 (WSM-R), and R21AI164578 (DJK). We thank Soumitra Guin and Marvin Meyers (St. Louis University) for synthesis and purification of CWHM-974.

